# Apolipoprotein A1 and high-density lipoprotein limit low-density lipoprotein transcytosis by binding scavenger receptor B1

**DOI:** 10.1101/2022.08.08.503162

**Authors:** Karen Y. Y. Fung, Tse Wing Winnie Ho, Zizhen Xu, Dante Neculai, Catherine A. A. Beauchemin, Warren L. Lee, Gregory D. Fairn

## Abstract

Atherosclerosis results from the deposition and oxidation of low-density lipoprotein (LDL) and immune cell infiltration in the sub-arterial space leading to arterial occlusion. Numerous studies have shown that transcytosis transports circulating LDL across endothelial cells lining the blood vessels. LDL transcytosis is initiated by binding to either Scavenger Receptor B1 (SR-B1) or Activin A receptor-like kinase 1 (ALK1) on the apical side of endothelial cells leading to its transit and release on the basolateral side. Individuals with elevated levels of circulating high-density lipoprotein (HDL) are partly protected from atherosclerosis due to its ability to remove excess cholesterol and act as an antioxidant. Apolipoprotein A1 (APOA1), an HDL constituent, can bind to SR-B1, raising the possibility that APOA1/HDL may also compete with LDL for SR-B1 binding and thereby limit LDL deposition in the sub-arterial space. To examine this possibility, we used *in vitro* approaches to quantify the internalization and transcytosis of fluorescent LDL in coronary endothelial cells. Using microscale thermophoresis and affinity capture, we find that SR-B1 and APOA1 directly interact, and that binding is enhanced when using the cardioprotective variant of APOA1 termed Milano (APOA1-Milano). In a murine model, transiently increasing the levels of HDL reduced the acute deposition of fluorescently labeled LDL in the atheroprone inner curvature of the aorta. Reduced LDL deposition was also observed when increasing circulating wild-type APOA1 or the APOA1-Milano variant, with a more robust inhibition from the APOA1-Milano. The results suggest that HDL may limit SR-B1-mediated LDL transcytosis and deposition, adding to the mechanisms by which it can act as an atheroprotective particle.

## Introduction

Heart attacks and strokes arise from the development of atherosclerosis, an inflammatory disease initiated by the accumulation of cholesterol-rich low-density lipoprotein (LDL) in the intima of the arterial wall following its transit across the endothelium. This is especially apparent in regions of the vasculature that experience disturbed, non-laminar blood flow.^1^ The excess intimal LDL is subject to oxidation, with the resulting oxidized LDL (oxLDL) contributing to the activation of the endothelium, leading to the accumulation of monocytes/macrophages that use scavenger receptors to internalize the excess lipoproteins.^2^ Unfortunately, macrophages overloaded with oxLDL-derived cholesterol differentiate into foam cells.^2^ Unlike macrophages, foam cells lose their ability to effectively clear dead cells and egress from the subendothelial space.^2^ Ultimately, foam cells die, which further contributes to the accumulation of necrotic cells.^2^ Over time, the collection of lipids, inflammatory cells, cellular debris, and foam cells is accompanied by smooth muscle cell infiltration into the intima to form a fibrous cap overlaying the necrotic core.^2,3^

The initial step in this process is the transit of LDL through the endothelial cell cytoplasm using the process of transcellular transport, commonly referred to as transcytosis.^4^ In this instance of transcytosis, LDL is internalized into vesicles at the apical surface and then transported through the cytoplasm and subsequently undergoes exocytosis and release at the basolateral surface. We and others have found that LDL transcytosis is mediated by its binding to the Scavenger Receptor B1 (SR-B1)^4,5^ and Activin receptor-like kinase 1 (Alk1).^6^ Similar to LDL, high-density lipoprotein (HDL) must also undergo transcytosis to move from the circulation to the intima of blood vessels.

HDL is cardioprotective, yet the exact mechanisms by which this occurs remain unclear and is likely multifactorial.^7^ The most highly ascribed function of HDL is as a cholesterol acceptor for reverse cholesterol transport, a process dependent on transcytosis to enter and exit the intimal space.^8^ However, HDL is also an anti-inflammatory particle^9–17^ that can promote endothelial integrity,^16,18–20^ ^12–15^ and induce the production of the vasodilator prostacyclin.^21^ Additionally, HDL carries microRNAs involved in regulating the vasculature, inflammation^22^, and SR-B1 expression.^23^ We and others have demonstrated that HDL transcytosis is also mediated by SR-B1^24,25^ and the ATP-binding Cassette (ABC) G1 transporters.^25^

SR-B1 was first identified by its ability to bind to modified LDL^26^ but has been since shown to bind various other ligands, including other lipoproteins,^26–28^ bacterial wall components and apoptotic cells.^29–34^ Commonly, SR-B1 is known as the HDL receptor with the apparent dissociation constant (K_d_) of the HDL-SR-B1 interaction being more robust than that of LDL, implying that HDL is the preferred SR-B1 ligand.^35^ As such, HDL can effectively outcompete LDL for binding to SR-B1, whereas LDL can only partially compete with HDL for SR-B1 binding.^36–38^ The amino acids on SR-B1 involved in binding with HDL partially overlap with those involved in LDL binding,^39^ suggesting that SR-B1 cannot simultaneously bind both particles. Since LDL and HDL both bind SR-B1, the ability of HDL to outcompete LDL for interaction with SR-B1 may allow HDL to regulate LDL transcytosis across the endothelium.

The ability of HDL to interact with SR-B1 is attributable to the main protein component, apolipoprotein A1 (APOA1),^40–43^ found exclusively in HDL. APOA1 is composed of several amphipathic helical motifs,^44^ required to maintain the structure of the HDL particle. APOA1 is also responsible for many of the beneficial properties of HDL. Although most plasma APOA1 exists as part of HDL in a lipidated state, a small transient pool of lipid-free APOA1^45^ exists to facilitate the exchange of APOA1 amongst different populations of HDL. Referred to as HDL-APOA1 Exchange (HAE), this process is reduced in samples from atherosclerosis patients and those with at least one cardiovascular disease (CVD) risk factor.^46^ Importantly, hypoalphalipoproteinemia, a disorder where APOA1 mutations prevent its synthesis, led to lowered plasma HDL levels and premature cardiovascular diseases.^47,48^ Together, this highlights the importance of APOA1 in lipid-free and lipid-associated forms in providing protective effects impeding cardiovascular pathogenesis. Furthermore, several naturally occurring APOA1 variants have been identified, some of which increase susceptibility to developing CVD while others offer protection. The APOA1-Milano variant has been of great interest in previous studies. This variant possesses an arginine 173 to cysteine substitution that facilitates the formation of an intermolecular disulfide bond between two APOA1 molecules.^49^ Curiously, carriers of the APOA1-Milano mutation have a variety of phenotypes, including ∼50% lower circulating plasma HDL and APOA1 levels, elevated triglycerides, and a change in HDL subtypes with elevated HDL_3_ and decreased HDL_2_,^50,51^ but exhibit a low incidence of CVD.^49,52^ How the ApoA1-Milano variant interacts with SR-B1 or if it can oppose LDL transcytosis is unknown.

Given that HDL and LDL bind to SR-BI, we hypothesized that competition for the receptor (and thus inhibition of LDL transcytosis) could be another mechanism by which HDL protects against atherosclerosis. We demonstrate that recombinant APOA1 antagonizes LDL internalization and transcytosis in human coronary artery endothelial cells (HCAECs). This was potentiated when APOA1 was reconstituted as a minimally lipidated HDL-like particle. The *in vitro* binding assays suggest APOA1-Milano is a better ligand for SR-B1 than the wild-type protein. In a murine model, acute increases in HDL, APOA1, and the Milano mutant all limit the deposition of LDL in the aorta. The results reveal an unappreciated mechanism by which APOA1/HDL is anti-atherogenic.

## Methods

### Cell Culture

Human coronary artery endothelial cells (HCAECs, Promocell – C-12221, lot 425Z019.1) were cultured in EGM2TM BulletKit^TM^ medium (Lonza Switzerland, CC-3162) and grown at 37°C at 5% CO_2_. Henrietta Lack cells (HeLa, ATCC) were cultured in Dulbecco’s Modified Eagle’s Medium (DMEM, Corning) supplemented with 10% FBS (Wisent) and grown in the same environmental conditions. For the HCAECs, the medium was changed every 2 to 3 days of culture and experiments were performed at least 2 days after seeding on 0.1% gelatin-coated 25 mm round coverslips.

### DNA Transfection

HeLa cells were transfected using X-tremeGENE^TM^ HP DNA transfection reagent (Sigma-Aldrich) at a ratio of 1:3 using 0.5 μg of DNA in 100 μL of Opti-MEM per 12 well. Cells were transfected 1 day after seeding, and internalization experiments were performed 1 day after transfection.

### Production of APOA1 Mutants

APOA1 variants were generated using the Phusion Site-Directed mutagenesis Kit (Thermo Fisher Scientific), E.coli codon-optimized cDNA encoding human APOA1^24^ and the following primers to generate the Milano variant: Forward: AACTGCGGCAATGCCTTGCAGC and Reverse: CGTCAGAGTACGGCGCGAGATGTG. Expression and Purification of the APOA1 variants were performed using ClearColi® BL21(DE3) (Lucigen).^24^ APOA1 was eluted from Talon® Metal Affinity Resin (Takara) using 0.1 M EDTA. Eluants were dialyzed with PBS at 4°C to remove 0.1 M EDTA and any residual Guanidine HCl; the dialysis buffer was changed twice before APOA1 protein collection. Purity was assessed by running dialyzed eluants on SDS-PAGE, and concentrations were calculated using the molar extinction coefficient of APOA1 at 280 nm and measuring the absorbance at 280 nm.

### Cloning and purification for microscale thermophoresis experiments

The full-length APOA1-WT, APOA1-Milano, and the N-terminal His_6_-tagged luminal domain of human SR-B1 (amino acid 33 – 444) were cloned into PB-T-PAF^53^ downstream and in frame with the protein A tag (protein A-APOA1, protein A-APOA1-Milano, and protein A-His_6_-*h*SR-B1^54^), by using a two-step PCR strategy with overlapping primers encompassing the desired mutation and sequence verified.

Recombinant APOA1, APOA1-Milano, and His-tagged SR-B1 ectodomain (His_6_.*h*SR-B1 (amino acid 33 - 444)) were expressed in 293F mammalian cells^53^ and purified as described previously.^54^ The protein concentrations were determined using a BCA assay.

### Generating reconstituted HDL particles containing APOA1 Variants

#### Incorporation of APOA1 to DMPC and DiI

Lipid-associated APOA1 were generated following previous protocols with slight modifications.^55,56^ Briefly, 1,2-dimyristoyl-sn-glycero-3-phosphocholine (DMPC) (Avanti Polar Lipids), with or without 1,1’ -Dioctadecyl-3,3,3’,3’-Tetramethylindocarbocyanine Perchlorate (DiI) (Sigma), were transferred to a 5 mL or 10 mL round bottom flasks manufactured from borosilicate 3.3 glass with a 14/20 Joint size (VWR) and dried to a film by streaming N_2_. This film was resuspended in methanol and dried to film by N_2_ streaming. The film was resuspended in 25 mM HEPES in PBS and sonicated at 24°C for 20 min or until the lipid film was fully resuspended. Recombinant APOA1 equilibrated to 24°C was added to the resuspended lipids, and further sonicated for 30 min at 24°C or until turbidity disappeared. This solution was incubated at 24°C overnight to allow APOA1 incorporation into vesicles.

#### Potassium Bromide Gradient Ultracentrifuge

Lipidated APOA1 was separated by buoyancy in a KBr gradient by ultracentrifugation. Briefly, the lipidated APOA1 was transferred to the bottom of a ½-inch × 2-inch ultracentrifuge tube and topped to 2 mL using PBS containing 0.3 mM EDTA. The density of this solution was adjusted to [2.45-2.5] g/mL using KBr. An additional 2 mL of 1.063 g/mL PBS containing 0.3 mM EDTA was layered on top. This was then centrifuged at 35,000 rpm for 24 h at 4°C in a Beckman Optima LE-80 Ultracentrifuge using a SW55Ti rotor with maximum acceleration and without breaking at the end of the spin. Next, 200 μL fractions were collected, and density was calculated by measuring the weight of the 200 μL fraction. Absorbances at 280 nm and 555 nm were measured using the NanoDrop^TM^ 2000/2000c (Thermo Scientific) to determine the presence of APOA1 and DiI (respectively) in each fraction. Fractions corresponding to the appropriate densities containing sufficient APOA1 were pooled and dialyzed to remove the dialysis.

#### Native Gel Electrophoresis of Lipoproteins

Samples were resolved using a 4-12% polyacrylamide gradient gel (pH 8.75) using the Tris-glycine buffer system (no SDS and non-reducing). The gel was first imaged to detect the DiI label and then stained with Coomassie to detect protein.

### Internalization Experiments

#### DiI-LDL Internalization

HCAECs were washed twice with cold Dulbecco’s Phosphate Buffered Saline supplemented with calcium and magnesium (PBS+, Sigma). 10 μg/mL DiI-LDL^4^ in the presence of PBS or the various unlabelled APOA1 proteins or rHDL^APOA1^ (of density 1.1 g/mL) in serum-free RPMI (Gibco, 1640) were incubated with HCAECs for 20 min at 37°C to allow for internalization. After 20 min, DiI-LDL was removed, and HCAECs were washed twice with PBS+ and fixed in 4% paraformaldehyde.

#### DiI-rHDL^APOA1^ Internalization

HeLa cells were washed twice with PBS+, and 5 μg/mL of DiI-labelled rHDL with different APOA1 variants (DiI-rHDL^APOA1^) in the presence of PBS or the various unlabelled APOA1 variant proteins in DMEM supplemented with 5% BSA were incubated with HeLa cells for 1 h at 37°C to allow for internalization. After 1 h, the DiI-rHDL^APOA1^ was removed, and HeLa cells were washed twice with PBS+ and fixed in 4% paraformaldehyde.

#### Microscopy

The Quorum Technologies (Guelph, Ontario) Diskcovery spinning disk confocal/TIRF system was used to image the DiI-LDL. This system is based on a Leica DMi8 with a Nipkow spinning disk with a pinhole size of 50 μm, a Hamamatsu ImageEMX2 EM-CCD camera, with diode-pumped solid-state lasers (405 nm, 488 nm, 561 nm, and 637 nm) and a 63x oil immersion objective with a 1.47 numerical aperture. For DiI-rHDL^APOA1^ internalization, a Quorum spinning disk confocal microscope (Yokogawa CSU-X1 spinning disk head with Borealis, Hamamatsu ImageEMX2 EM-CCD camera at 63x objective, numerical aperture 1.4, with diode-pumped solid-state lasers 405 nm, 488 nm, 552 nm, and 730 nm) was used. In both experiments, z-slices encompassing the whole cell were acquired at intervals of 0.2 μm.

For the HDL experiments (**Figure 5**), the entire aortic arch was imaged by automated tiling with 16 z-slices, spanning 15 μm in depth, per field of view using a ZEISS LSM 900 microscope with a 20x/0.8 NA objective, and 405 nm, 488 nm, 561 nm laser lines acquired using the GaAsP-PMT detectors. The same approach was used for the APOA1 experiments (**Figure 6**) with the exception that the microscope used was a ZEISS LSM 700 with a 20x/0.8 NA objective, 405 nm, 488 nm, 555 nm, laser lines, and acquired with a PMT detector. In all experiments, acquisition settings, including laser power and scan speed, were kept constant between conditions to directly compare the intensity data for samples processed on the same day. All acquisitions were acquired with ZEN Black software. In each experiment, one animal that did not receive AF568-LDL was processed to determine the background (tissue autofluorescence). To quantify LDL deposition, ImageJ was used to subtract the background fluorescence, and integrated densities of the Alexa Fluor 568-LDL were quantified and normalized to the (CD31-positive) aortic area.

#### Analysis and Quantification

The amount of DiI-LDL and DiI-rHDL^APOA1^ internalization per cell was calculated by the sum of the raw integrated density of the DiI channel within the region of interest (the cell) across all z-slices measured using Fiji (NIH). Due to the varying sizes of HeLa cells in the DiI-rHDL^APOA1^ experiments, which may affect the absolute total DiI-rHDL^APOA1^ fluorescence, the summed raw integrated density of DiI-APOA1 was further normalized by dividing by the volume of the cell which was calculated by multiplying the area of the cell (the region of interest from which the raw integrated density was measured from as determined by the GFP channel) by the number of slices analyzed. Values plotted for DiI-LDL internalization are the absolute DiI-LDL fluorescence per cell analyzed. Values plotted for DiI-rHDL^APOA1^ internalization are normalized to the DiI-rHDL^APOA1^ uptake seen in the GFP transfected alone control.

### In vitro APOA1-SR-B1 Binding Assay

The interaction between SR-B1 and APOA1 or rHDL was assessed through an *in vitro* binding assay using GFP nanotrap beads. The GFP nanotrap was made as previously.^57^ For the binding assay, HeLa cells overexpressing GFP-SR-B1 (as a fusion protein) or GFP alone (control) were lysed in lysis buffer (0.5% NP-40, 10 mM glycine, protease inhibitor in PBS). The soluble fraction of the lysates (supernatant after centrifugation for 13,000 rpm for 10 min at 4°C) were incubated with pre-equilibrated GFP nanotrap beads overnight with end-to-end rotation at 4°C to immobilize GFP or GFP-SR-B1 onto beads. The beads were washed 3x with lysis buffer, after which the APOA1 variant protein was added and topped with PBS to a final total volume of 1 mL. This reaction was further incubated with end-to-end rotation at 4°C for 3 h to allow for the APOA1 interaction with GFP or GFP-SR-B1. The GFP or GFP-SR-B1 beads were then washed 3x - 4x with high salt (1.5 M) PBS to remove non-specific interaction, and SDS-PAGE loading buffer (either in the presence or absence of DTT, where indicated) was added to elute remaining proteins off the beads. The sample was analyzed through western blot and immunoblot with antibodies against APOA1 (Abcam, clone [EP1368Y]) and GFP (Bio X Cell, clone [F56-6A1.2.3]) to assess the amount of APOA1 bound and the amount of GFP or GFP-SR-B1 immobilized onto the beads between different conditions. For quantification of the APOA1-SR-B1 interaction, the intensity of all the bands were measured through BIO-RAD Image Lab 5.2.1. The APOA1 band intensity, when incubated with SR-B1-GFP, was first divided by the band intensity of SR-B1-GFP as minimally variable amounts of immobilized SR-B1-GFP were observed per condition. This value was then divided by the intensity of the corresponding APOA1 band in the input lanes to account for slight differences in the amount of APOA1 added to each condition. Lastly, Milano values were normalized to WT values and this was plotted to visualize the differences between the two variants better.

### Microscale thermophoresis (MST)

MST experiments were performed on a Monolith NT.115 instrument (NanoTemper Technologies). Purified His-tagged SR-B1 protein (His_6_.*h*SR-B1(amino acids 33 - 444), was labeled using the Monolith His-Tag Labeling Kit Red-tris-NTA (NanoTemper Technologies) and diluted in MST buffer (NanoTemper Technologies) to a concentration of 50 nM, was mixed with APOA1 or APOA1-Milano in MST buffer (PBS with 0.05%Tween) at indicated concentrations ranging from 1 nM to 8 μM at room temperature. Fluorescence was determined in a thermal gradient at 40% LED power and high MST power generated by a Monolith NT.115 from NanoTemper Technologies. Data were analyzed using Affinity Analysis v2.3.

### Measuring Transcytosis Using Total Internal Reflection Fluorescence Microscopy

Total internal reflection fluorescence (TIRF) microscopy was conducted using live HCAEC monolayers seeded on glass coverslips and placed in a chamblide imaging chamber. Videos were taken using the Quorum Diskcovery TIRF system, where 150 images were taken at 100 ms exposure with a penetration depth of the illumination of 100 nm into the basal membrane.

Confluent monolayers were pre-chilled for 5 min in cold HEPES-buffered RMPI (HPMI) media at 4°C. HPMI media was removed, and 20 μg/mL DiI-LDL^4^ with the recombinant APOA1 variant or rHDL^APOA1^ or PBS in cold HPMI was added and incubated for 10 min at 4°C to allow binding. Cells were then washed twice with cold PBS+ to remove unbound ligand, and prewarmed HMPI was added, after which the cells were placed on the live imaging stage. After approximately 1 min 45 s, imaging of the cell began. Cells in confluent regions, identified through the number of nuclei in the DAPI channel stained by Nucblue^TM^ Live cell staining (Thermo Fisher Scientific) added at the membrane-binding step, were imaged. A total of 10 fields of view were imaged per condition, where videos acquired were staggered by about 1 min.

The number of exocytosis events was calculated using a previously developed single-particle-tracking algorithm^4,58^ with modifications.^59^ Briefly, the local background signal was subtracted, and images were thresholded to create a binary image (mask). Putative vesicles were identified based on size (60 nm to 200 nm in diameter), circularity (>0.2), and intensity (threshold of 15% above mean image intensity). Vesicles were tracked based on maximum probability, which was dependent on the change in centroid particle and change in peak intensity. Exocytosis involves vesicles docking at the PM before membrane fusion and cargo release; hence the identification of transcytosis events was limited to vesicles that were stationary (potentially docked). Vesicle diffusivity (mean-squared displacement) was quantified, and those exhibiting subdiffusive behaviour before exocytosis (α<1) were included. From this subdiffusive population, vesicles that underwent a reduction of at least 3 standard deviations in fluorescent intensity during the last 5 time points of their tracks and did not reappear for 5 subsequent frames were counted as exocytosis events.

### LDL Perfusion of Mouse Aorta

The acute accumulation of LDL in the inner curvature of the murine aorta was measured as previously reported.^58^ Briefly, male C57BL/6J mice (Jackson Laboratory) were housed in a temperature-controlled room on a standard 12 h light-dark cycle, with *ad libitum* access to water and food. All experiments were conducted during the light phase of the cycle, between 10 weeks to 13 weeks of age. All experimental procedures were approved by St. Michael’s Hospital Research Ethics Board (19-314) and Animal Care Committee (ACC 899 and 204) and were in accordance with the guidelines established by the Canadian Council on Animal Care.

For the HDL competition experiments, mice were anesthetized and retro-orbitally injected with or without 1.25 mg of HDL at the same time as 200 µg of Alexa Fluor 568 conjugated-LDL (AF568-LDL). In APOA1 and APOA1-Milano competition experiments, mice were anesthetized and administered saline, 0.4 mg ApoA1, or 0.4 mg ApoA1 Milano by retro-orbital injection. Thirty minutes later, mice were administered 200 µg of AF568-LDL via retro-orbital injection. In both experiments, 30 minutes post-injection of AF568-LDL, mice were euthanized by exsanguination, and the aortas were slowly perfused with 5 ml of PBS containing calcium and magnesium (PBS^+/+^) followed by 5 ml of 10% buffered formalin through the left ventricle. Hearts were dissected and fixed with 10% buffered formalin for 60 minutes. After fixation, adventitial fat and tissue on the aortic arch were removed using microdissection scissors. The aortic arch was removed from the descending aorta, aortic branches, and aortic root.

Isolated aortas were permeabilized with 0.5 % Triton X-100 and 10 % DMSO in PBS^+/+^ for 5 minutes. After washing, aortas were incubated with 10 µg/ml rabbit anti-mouse CD31 primary antibody (NB100-2284; Novus) in 5% BSA at 4 °C overnight. Aortas were then washed and incubated with 7.5 µg/ml anti-rabbit Alexa Fluor 488 secondary antibody (Jackson ImmunoResearch) in 5 % BSA in room temperature for 1 hour followed by 2.5 ug/ml DAPI (Roche) for 20 minutes. After washing, aortas were mounted *en face* with Dako fluorescence mounting medium (Agilent). Samples were examined using laser scanning confocal microscopy as described in the ‘Microscopy’ section.

### Statistics and Graphic Representation

Each experiment was repeated n times and within each experiment (each k out of n, where k represents each experimental replicate), where each n represents an experiment done at a different time and using a different passage of cells. The fluorescence of several individual cells was measured either with cells alone (**blank**) or in the presence of a fluorescently tagged compound (e.g. DiI-LDL) to which either **control** is added (e.g. PBS) or a **condition** is investigated (e.g. addition of an APOA1 variant).

For LDL transcytosis experiments using recombinant APOA1 as competitors, LDL internalization experiments using rHDL^APOA1^ as competitors, rHDL^APOA1^internalization experiments using recombinant APOA1 as competitors, and the internalization of DiI-DMPC particles that lack APOA1 experiments, the graphs were generated using GraphPad Prism 5.0 (GraphPad Software Inc., La Jolla, CA, USA), where data were presented as averages per experiment ± SEM. The statistics were performed using Student’s t-test or ANOVA (GraphPad Prism 5.0; GraphPad Software Inc., La Jolla, CA, USA) with Bonferroni’s or Dunn’s selected *Post-hoc* for raw or normalized data (respectively).

For the LDL internalization experiments and LDL transcytosis experiments using rHDL^APOA1^ as competitors, the graphs were generated using python’s matplotlib library. The change in fluorescence for a given condition, relative to the control (ctrl), for experiment replicates k, was computed as:

> Change_ctrl_k = { [ ctrl_i – mean(blank) ] / [ ctrl_j – mean(blank) ] } for all i,j

> Change_cond_k = { [ cond_i – mean(blank) ] / [ ctrl_j – mean(blank) ] } for all i,j

Where: cond_i is the fluorescence for each cell under the condition (i.e. I from 1 to Ncells_cond), ctrl_j is the fluorescence for each cell in control (i.e. j from 1 to Ncells_ctrl), and mean(blank) is the average fluorescence for all cells alone.

As such, Change_cond_k is a set of fluorescence measures for experiment replicate k, represented as dots of the same colours in such figures, and there are Nchange_k = Ncells_cond * Ncells_ctrl, such measures in each replicate k of the n total, replicates conducted for each control or condition. Because steps in Change_cond_k are relative to both blank and control measurements, the conditions within a particular experiment are thus subtracted so the set of n replicates of Change_cond_k sets can be combined as a single set, Change_cond, which contains Nall = Nchange_k * n points. The bars appearing next to the dots correspond to the edge of the 95^th^ (dark grey) and 68^th^ (light grey) percentiles and the median (black horizontal bar) for the Change_cond set of measurements.

The fold-change and p-value between the control and the condition are computed by taking each of the Nall_cond points in Change_cond and dividing them by each of the Nall_ctrl points in Change_ctrl. The median and 95^th^ percentile of this set of Nall_cond * Nall_ctrl points are reported in the text. If the condition corresponds to an increase in fluorescence, i.e. if the median fold-change is greater than one, the p-value is the probability of the null hypothesis, i.e. that the condition does not correspond to an increase in fluorescence (fold-change equal to or less than one). This is computed as the Binomial likelihood of get f_fail times n_trial failures out of n_trial attempts. Here f_fail is the fraction of the Nall_cond times Nall_ctrl points where the fold-change was equal to or less than one, and n_trial is the number of independent points which is not Nall_cond * Nall_ctrl. Indeed, while set Change_ctrl contains Ncells_cond times Ncells_ctrl measures, these measures are not completely independent in that each cond_j is re-used Ncells_ctrl times, i.e. divided by all ctrl_j measures. To correct for this, we take n_trial as the sum of the number of cells whose fluorescence was measured under the condition for each replicate, i.e. n_trial is the sum of Ncells_cond for each of the n replicates.

For the other experiments, a student’s t-test was utilized. For LDL perfusion in vivo experiments, 1-way ANOVA was performed using GraphPad Prism software, version 9.0 (La Jolla, CA, USA). Individual comparisons between samples were analyzed using Šidák post hoc test and data were presented as mean per experiment ± SEM.

## Results

### Inhibition of LDL Transit Across the Endothelium by Excess Recombinant Wild-Type APOA1

To address whether wild-type APOA1 (APOA1-WT) can inhibit LDL transit across endothelial cells, we expressed and purified recombinant APOA1 from ClearColi^TM^, an E.coli mutant deficient in lipopolysaccharide (**Supplemental Figure 1**).^24^ Firstly, we examined the effect of unlabelled APOA1on DiI-LDL internalization in HCAECs. After incubation of DiI-LDL and unlabelled APOA1 with HCAEC for 15-20 min at 37°C, cells were washed, fixed, imaged using confocal microscopy and subsequently analyzed using ImageJ to measure the amount of LDL internalization by quantifying the fluorescence intensity of DiI (**Figure 1A**). The internalization of DiI-LDL by the primary HCAECs (**Supplemental Figure 2A**) exhibited high variability, likely due to the culturing of the primary human cells as such cells displayed 10^4^ to 10^6^ fluorescence units (total integrated fluorescence per cell) (**Supplemental Figure 2B**). This variability made the assay less sensitive in detecting subtle changes in DiI-LDL internalization. Hence the DiI-LDL fluorescence signal of each cell per condition was compared to each cell in control (PBS alone), and the median change was compared between treatments. As a result, a 20-fold excess (by protein mass) of unlabelled APOA1 led to a significant (p-value = 10^−6^) 1.96-fold reduction (median fold change of 0.51 or log2(−0.97)) [95% CI: (0.035, 150.4)] in the median DiI-LDL internalization compared to the PBS treated control cells (**Figure 1B**). This suggests that APOA1 inhibits the internalization of DiI-LDL in HCAECs.

**Figure 1.**
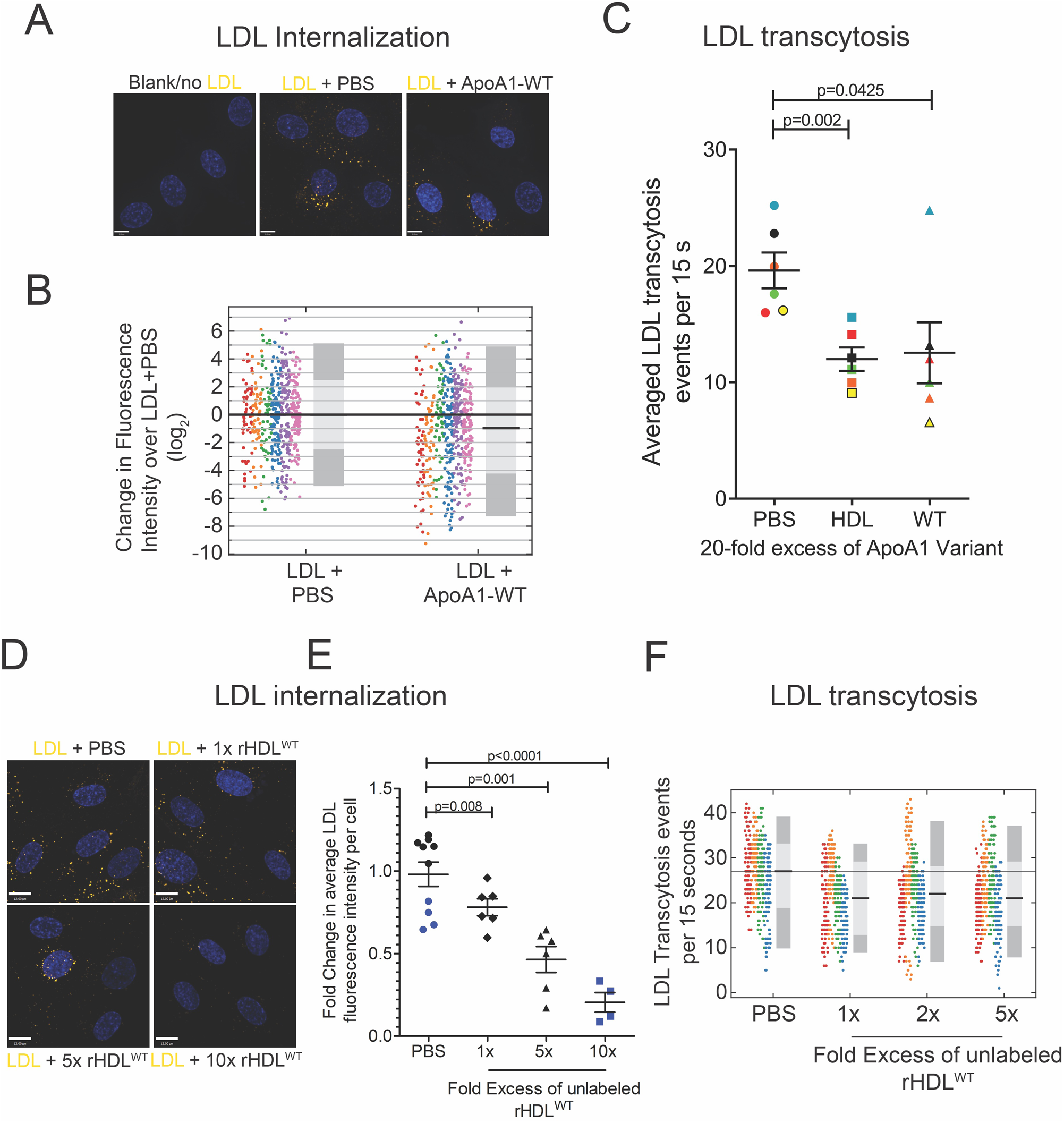
Unlabeled APOA1/reconstituted HDL inhibits DiI-LDL transcytosis in human coronary endothelial cells. **(A)** Representative images of internalized DiI-LDL in HCAECs treated with 20-fold excess (by protein mass) unlabeled APOA1-WT. **(B)** Quantification of fold change in DiI-LDL signal per cell normalized to the PBS control (DiI-LDL alone) along with median fold change presented as the black bar and the 95% (dark grey) and 68% (light grey) confidence interval. Replicates are color-coded, n = 6 repetitions, where at least 40 cells were quantified per condition per repetition. **(C)** Inhibition of DiI-LDL transcytosis by 20x excess (by protein mass) unlabeled commercially purchased human purified HDL and also by 20x excess (by protein mass) unlabeled APOA1-WT as measured using TIRF microscopy. Presented are the average transcytosis events over a span of 15 secs per field of view after subtraction from a no DiI-LDL coverslip over n = 6 repetitions. Replicates are color-coded, where 10 cells were imaged per condition per repetition. **(D)** Representative images of DiI-LDL internalization competition by 10-, 5-, 1-fold excess (by protein mass) of rHDL^WT^ relative to DiI-LDL by protein mass. (**E**) Quantification of the amount of DiI-LDL uptake expressed as fold change compared to DiI-LDL added alone. Replicates are color-coded, n=5 for 1x & 5x and n=4 for 10x, where 60 cells were imaged and analyzed per condition per n. (**F**) Inhibition of DiI-LDL transcytosis by excess unlabeled rHDL^WT^ as measured using TIRF microscopy. Presented are the transcytosis events per cell over a span of 15 seconds per field of view after subtraction from a no DiI-LDL coverslip, along with average fold change presented as the black bar and the 95% (dark grey) and 68% (light grey) confidence interval. Replicates are color-coded, n=6, where 10 cells were imaged per condition per n.

To determine whether the inhibition of LDL internalization impacts transcytosis, we monitored the appearance of DiI-LDL in the TIRF plane of confluent monolayers. The presence of a 20-fold excess (by protein mass) of unlabelled APOA1-WT or commercially purchased human-isolated HDL with DiI-LDL during the receptor-binding step performed in the cold with HCAECs led to a significant decrease in DiI-LDL transcytosis by approximately 40% compared to DiI-LDL alone (PBS control) (p-value for ApoA1-WT < 0.05, the p-value for commercially purchased human isolated HDL <0.01) (**Figure 1C**). Together, these results demonstrate that APOA1 can cause a partial reduction of LDL internalization and that the attenuated uptake correlates with less release of LDL by transcytosis.

### Reconstituted HDL is a Better Competitive Inhibitor than Free-APOA1

Since lipid-free, recombinant APOA1 shows a weaker SR-B1 association^56^ we generated reconstituted APOA1 (rHDL^APOA1^) and examined its ability to inhibit LDL uptake and transcytosis. APOA1-WT was incorporated into unlabelled HDL-like particles consisting of 1,2-dimyristoyl-sn-glycero-3-phosphocholine (DMPC). Characterization of particle size and density of the resulting APOA1-WT particles (rHDL^WT^) indicated similar properties with native human HDL (**Supplemental Figure 3**). Due to the preferred association of SR-B1 with HDL_2_ compared to HDL_3_,^60^ rHDL^APOA1^with densities similar to that of HDL_2_ were used in subsequent experiments.

The ability of rHDL^WT^ to inhibit LDL internalization was examined in a dose-dependent manner. Various concentrations of rHDL^WT^ (10-, 5-, 1-fold excess relative to LDL by protein mass) were tested, noting that a 1-to 2-fold excess (by protein mass) represents the physiological ratio of HDL to LDL in healthy individuals. As expected, a dose dependence was observed where the concentration of rHDL^WT^ correlated with the percent inhibition of LDL internalization. Treatment with 10-fold excess (by protein mass) rHDL^WT^ reduced the LDL internalization by ∼72% (p-value <0.0001), whereas about a 50% decrease was seen at 5-fold excess (by protein mass) rHDL^WT^ (p-value = 0.001) followed by a 25% decrease with 1-fold excess (by protein mass) (p-value = 0.008) (**Figures 1D, E**). This suggests that at physiologically relevant concentrations, 1-to 2-fold excess (by protein mass) of rHDL^WT^, HDL trends towards partial inhibition of LDL internalization by HCAECs. Given the interest in SR-B1-mediated transcytosis, rHDL^WT^ was tested for its ability to inhibit DiI-LDL transcytosis using TIRF microscopy. In this assay, co-incubation of the various doses of rHDL^WT^ partially inhibited DiI-LDL transcytosis (1x = 20.7%, p-value = 0.001; 2x = 18.5%, p-value = 0.003; 5x = 19.2%, p-value = 0.003). Together, these results demonstrate that for physiological ratios of HDL to LDL, LDL uptake and transcytosis are attenuated (**Figure 1F**).

### APOA1-Milano is a better SR-B1 ligand than APOA1-WT

APOA1-Milano patients are protected from atherosclerosis, yet the mechanism for this protection is unclear. Since APOA1-Milano can form a homodimer and SR-B1 can multimerize,^61^ an increase in avidity could mean that APOA1-Milano and the resulting HDL are better ligands for SR-B1. If so, this raises the possibility that APOA1-Milano / Milano HDL is a better inhibitor of LDL transcytosis. Indeed, the low circulating HDL levels in APOA1-Milano carriers could result from enhanced SR-B1 mediated transcytosis of Milano HDL out of the bloodstream. To examine this possibility, we generated fluorescently labeled rHDL^WT^ and rHDL^Mil^ (DiI-rHDL^WT^ and DiI-rHDL^Mil^, respectively) by including DiI during particle formation. Next, we used these particles to monitor SR-B1-mediated internalization by confocal microscopy. HeLa cells transfected with a bicistronic vector encoding unlabelled SR-B1 and a soluble GFP (**Supplemental Figure 4A, B**) were used to measure the uptake of DiI-HDL. Compared to HeLa cells expressing soluble GFP or those expressing GFP-Alk1, the SR-B1 expressing cells displayed an almost 10-fold increase in fluorescence when incubated with DiI-rHDL^wt^ and close to 15-fold for DiI-rHDL^Mil^ (**Figure 2A, B**). As a control, DiI-containing DMPC liposomes (DiI-DMPC) devoid of APOA1 showed minimal uptake by SR-B1 expressing Hela cells (**Supplemental Figure 4C**).

**Figure 2.**
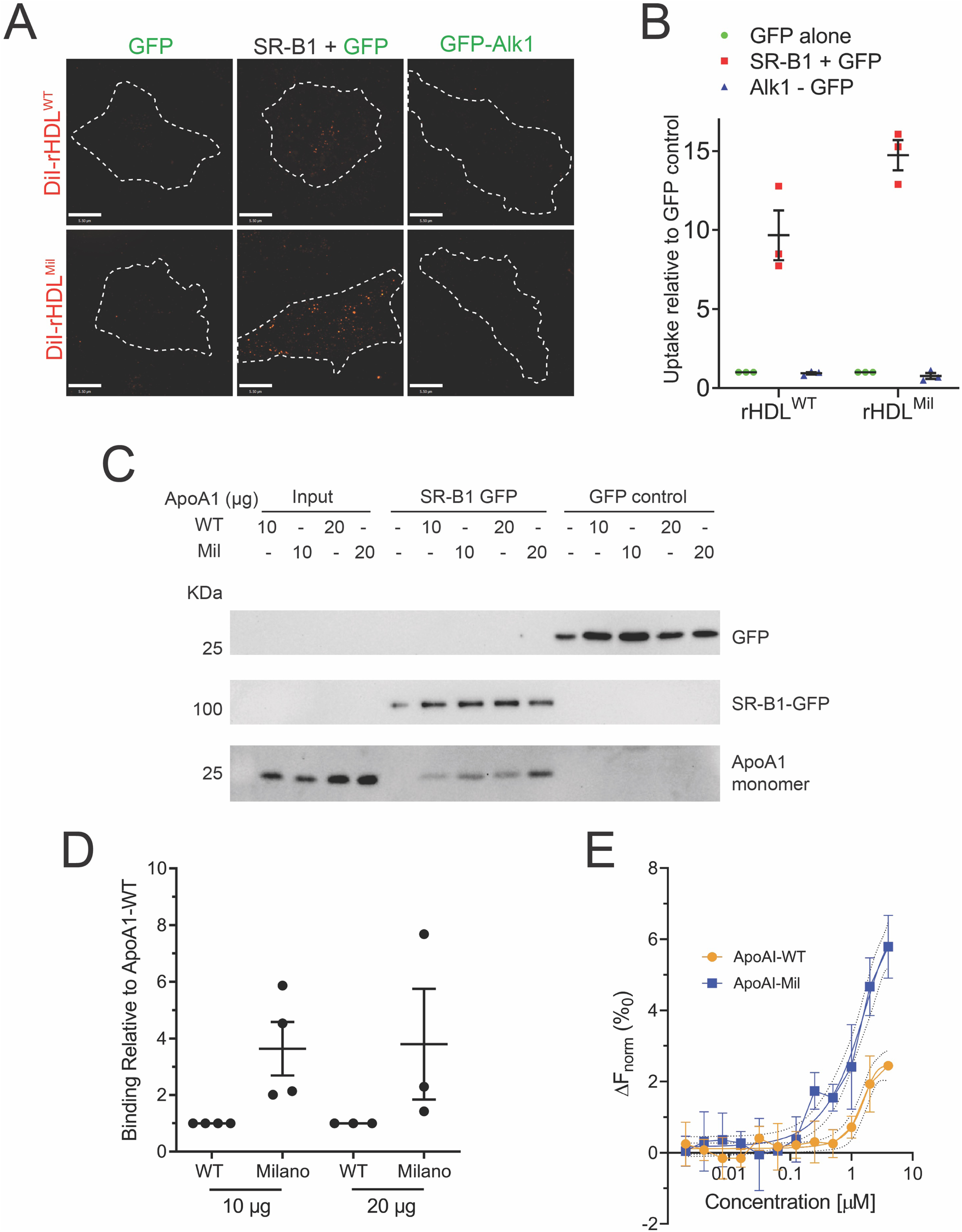
APOA1/reconstituted HDL interacts with SR-B1 while APOA1-Milano shows enhance binding to SR-B1 compared to APOA1-WT. **(A)** Representative images of rHDL^WT^ or rHDL^Mil^ incubated with HeLa cells transiently overexpressing SR-B1 + GFP, GFP-Alk1 and the control GFP alone. The dotted lines demarcate the boundary of the transfected cell (identified by GFP expression on the GFP channel) from neighboring cells. **(B)** The amount of rHDL^WT^ or rHDL^Mil^ signal per cell was averaged and normalized to the basal amount seen when transfected with GFP alone. n=3 for either experiment where 15 cells were imaged and analyzed per condition per n. **(C)** Representative western blot of the *in vitro* APOA1-SR-B1 binding assay using purified APOA1 variants, SR-B1-GFP fusion protein, and GFP along with the quantification **(D)** expressed as fold change compared to APOA1-WT interaction with SR-B1-GFP (fusion protein) that has been normalized to APOA1 in the input and amount of SR-B1 immobilized. n = 4 for 10ug APOA1; n = 3 for 20ug APOA1. **(E)** Binding of various concentrations of APOA1-WT or APOA1-Mil to 50nM SR-B1 as measured by microscale thermophoresis.

Next, we sought to more directly examine the ability of APOA1 to interact with SR-B1. To test this experimentally, we first performed an *in vitro* binding assay by immobilizing SR-B1-GFP overexpressed in HeLa cells onto GFP binding nanotrap beads, followed by incubation with recombinant APOA1-WT or APOA1-Milano. Association with SR-B1 was assessed by immunoblotting with anti-APOA1 antibody to visualize the amount of APOA1 protein that remains immobilized in the beads (interacting with GFP-SRB1). As shown in **Figure 2C, D**, APOA1-WT binding to SR-B1-GFP was detected on the western blot through the presence of the APOA1 band around 25 kDa. In parallel, the binding of APOA1-Milano to the immobilized SR-B1 was also analyzed. As shown in **Figure 2C, D**, 2-3-fold more APOA1-Milano was captured with SR-B1-GFP when normalized to input. As a control, APOA1-WT and APOA1-Milano were also incubated with GFP immobilized onto the GFP binding nanotrap beads, which resulted in no APOA1 bands, indicating the specificity of binding.

Next, to complement the pulldown assays, we determined the binding affinities of the APOA1-WT and APOA1-Milano to the ectodomain domain of SR-B1. To do this, we used microscale thermophore (MST)^62^ and examined the interaction between the recombinant His6X-tagged ectodomain of human SR-B1 (His_6_.*h*SR-B^33−444^) and full-length APOA1 and APOA1-Milano. These experiments generated the apparent Kd values to be 6.87 μM ± 0.23 μM for APOA1-WT and 2.06 μM ± 0.46 μM for APOA1-Milano, respectively (Fig 2E).

### Recombinant APOA1-Milano inhibits LDL internalization and transcytosis

Since APOA1-WT inhibits the uptake and transcytosis of LDL (Figure 1), we next examined if this ability is also present in the APOA1-Milano. First, DiI-LDL endocytosis was monitored in the HCAECs incubated with APOA1-WT or APOA1-Milano and compared to control cells treated with PBS. As shown in **Figure 3A, B**, adding 20-fold excess (by protein mass) of both APOA1 molecules decreased uptake. Specifically, the presence of 20-fold excess (by protein mass) of unlabelled recombinant APOA1-Milano also led to a significant (p-value = 10^−12^) 2.9-fold median reduction (median fold change of 0.347 or log2(−1.53)) [95% CI (0.05, 326.8)] in DiI-LDL fluorescence compared to DiI-LDL alone (PBS), indicative of a decrease in DiI-LDL internalization in HCAECs. Compared to APOA1-WT, APOA1-Milano appeared to be a slightly better inhibitor of LDL internalization (p-value < 0.05). Additionally, we find that APOA1-Milano can attenuate LDL transcytosis by ∼40% (p-value = 0.0085) (**Figure 3C**). Together, these results suggest that the Milano mutation does not impact the ability of APOA1 to compete for LDL internalization.

**Figure 3.**
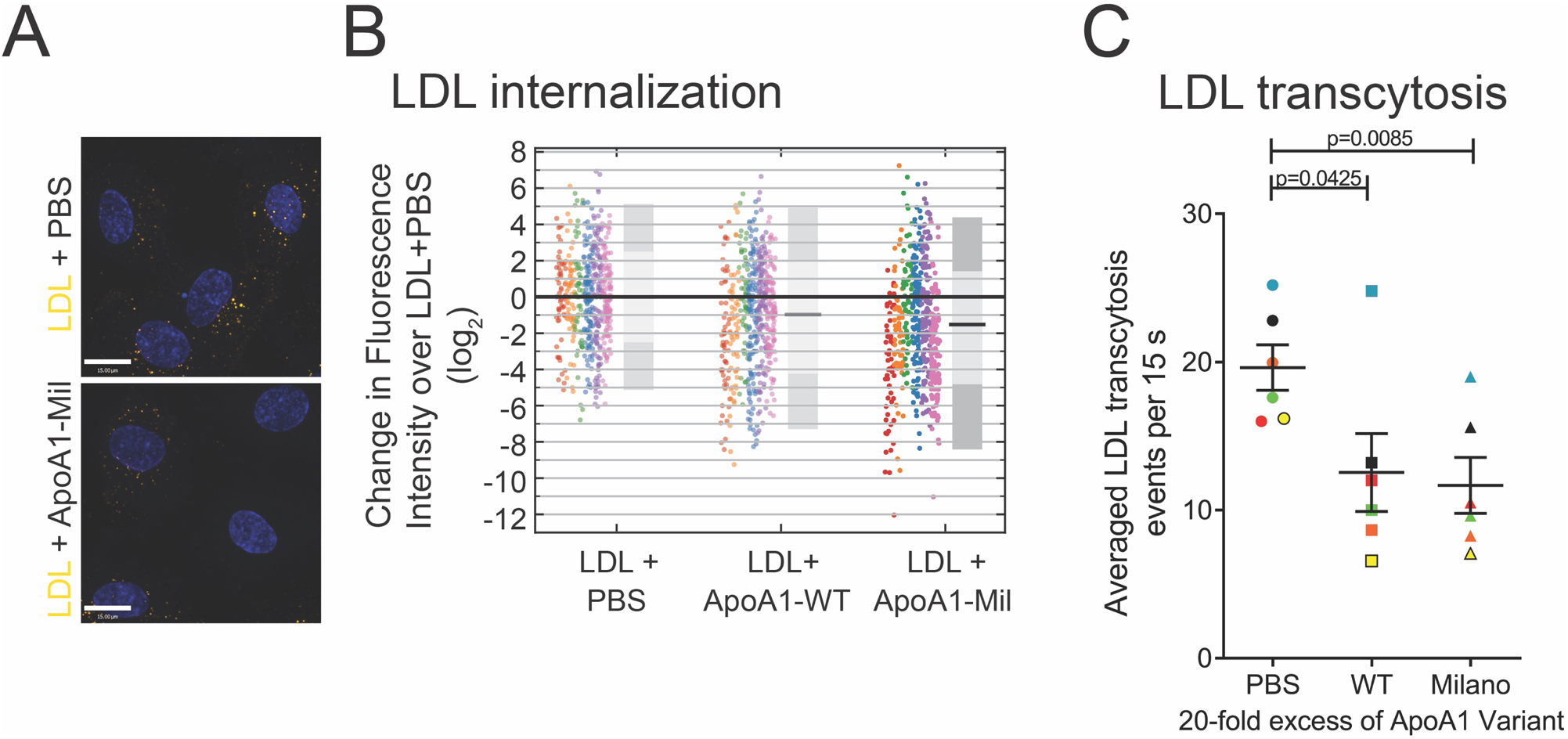
Recombinant APOA1-Milano inhibits DiI-LDL internalization and transcytosis in human coronary endothelial cells. The effect of excess APOA1-Milano on LDL uptake and LDL transcytosis were tested in the same experiment as APOA1-WT (Figure 1), hence repetition of the PBS control and APOA1-WT. **(A)** Representative images and the quantification of the impact of adding excess unlabeled APOA1-Milano on internalization of DiI-LDL in coronary endothelial cells. **(B)** Quantification of fold change in DiI-LDL signal per cell normalized to the PBS control (DiI-LDL alone) along with median fold change presented as the black bar and the 95% (dark grey) and 68% (light grey) confidence interval. Replicates are color-coded, n = 6 repetitions, where at least 40 cells were quantified per condition per repetition. **(C)** Inhibition of DiI-LDL transcytosis by excess unlabeled APOA1-Mil as measured using TIRF microscopy. Presented are the average transcytosis events over a span of 15 secs per field of view after subtraction from a no DiI-LDL coverslip n = 6. Replicates are color coded, where 10 cells were imaged per condition per n.

As previously noted, APOA1, when present in HDL particles, is a better ligand for SR-B1. Thus, we generated rHDL containing APOA1-Milano and assessed its ability to block LDL internalization and transcytosis by HCAECs ranging from physiological concentrations to supraphysiological levels up to 10-fold higher. As displayed in **Figure 4A** and quantified in **Figure 4B**, a physiological (1x) amount of rHDL^Mil^ reduced LDL uptake by ∼40% (p-value = 0.02), while 5x in rHDL^Mil^ led to inhibition of LDL uptake by ∼50% (p-value = 0.0002) and 10x showed inhibition of ∼60% (p-value = 0.0006). This degree of inhibition is comparable to siRNA-mediated silencing of SR-B1.^4^ This suggests that under physiological conditions, rHDL^Mil^ may prevent LDL internalization dependent on the SR-B1 receptor but has minimal impact on the ALK1-mediated internalization. Furthermore, the degree of inhibition caused by the rHDL^Mil^ at physiological concentrations was comparable, with a slight tendency towards being greater, than that obtained with the wild type containing rHDL (**Figure 1D**). Additionally, the impaired internalization of DiI-LDL translates to a ∼22.6% decrease in LDL transcytosis at physiological conditions (or 1x excess, p-value = 0.001) with minimal additional impact when supraphysiological amounts of HDL^Mil^ was used (2x = 11.5% decrease, p-value = 0.04; 5x = 21.1% decrease, p-value = 0.003) (**Figure 4C**). Together, these results demonstrate that Milano containing HDL can inhibit LDL uptake and transcytosis.

**Figure 4.**
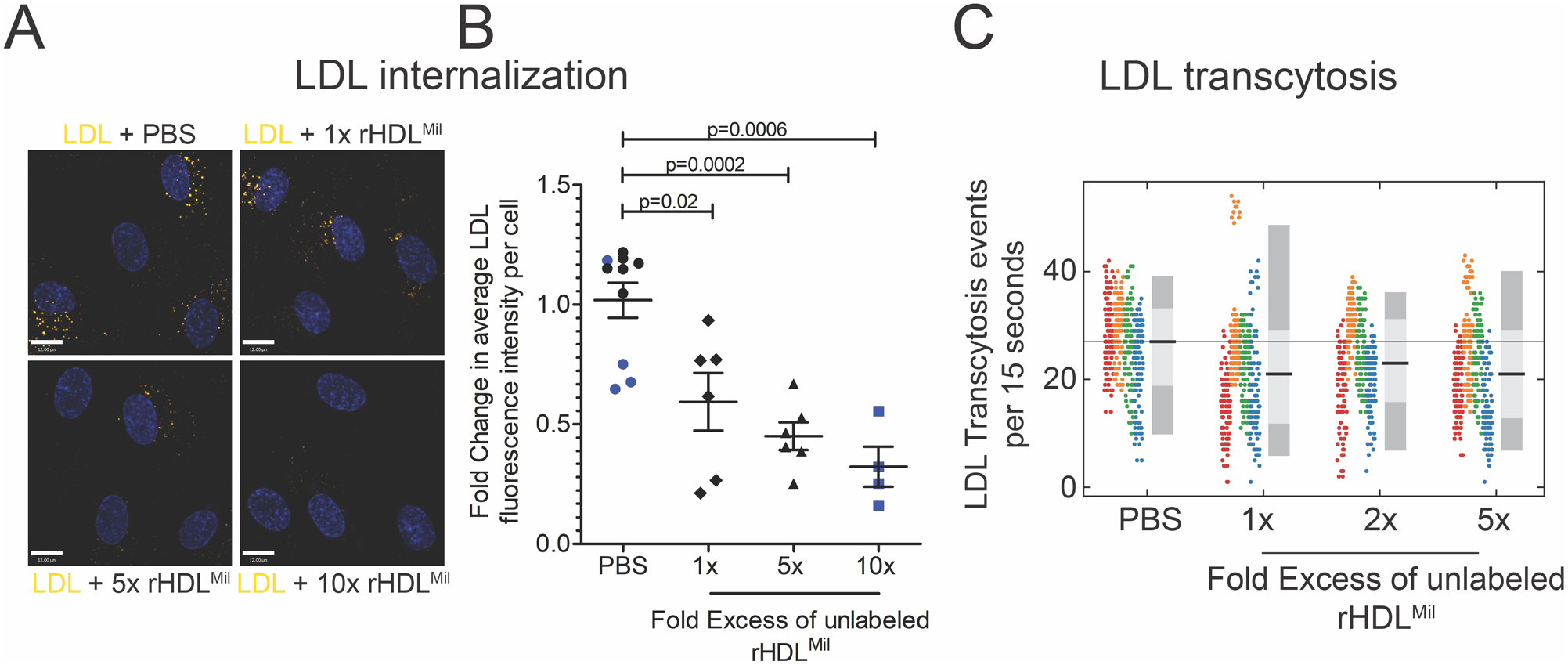
Inhibition of DiI-LDL internalization and transcytosis using physiological doses of unlabeled rHDL^Mil^ in human coronary endothelial cells. The effect of excess rHDL^Mil^ on LDL uptake and LDL transcytosis was tested in the same experiment as rHDLWT (Figure 1), hence the repetition of the PBS control. **(A)** Representative images of DiI-LDL internalization competition by 10-, 5-, 1-fold excess (by protein mass) of rHDL^Mil^ relative to DiI-LDL by protein mass. **(B)** Quantification of the amount of DiI-LDL uptake expressed as fold change compared to DiI-LDL added alone. Replicates are color-coded, n = 5 for 1x & 5x and n = 4 for 10x, where 60 cells were imaged and analyzed per condition per n. (**C**) Inhibition of DiI-LDL transcytosis by excess unlabeled rHDL^Mil^ as measured using TIRF microscopy. Presented are the transcytosis events per cell over a span of 15 secs per field of view after subtraction from a no DiI-LDL coverslip along with average fold change presented as the black bar and the 95% (dark grey) and 68% (light grey) confidence interval. Replicates are color-coded, n=6, where 10 cells were imaged per condition per n.

### The effects of APOA1 Milano on LDL deposition *In vivo*

We next sought to determine if a transient increase in HDL can inhibit the acute deposition of LDL in the aorta of male mice. For this experiment, we used freshly isolated HDL from healthy donors as previously^63^. The circulating levels of HDL-cholesterol in inbred mice strains range from 35 mg/dl up to 172 mg/dl, with C57/B6 having a value of ≈60 mg/dl.^64^ Thus, we co-administered via retro-orbital injection 1.25 mg of HDL-cholesterol, or saline as a control, to increase circulating HDL ∼2-fold, together with the Alexa Fluor 568-LDL (AF568-LDL). A graphical representation of the workflow is included in **Figure 5A**. Mice were euthanized 30 min post-injection, and the animals were processed as described in the Methods. Briefly, the aortic arch was isolated and stained with an anti-CD31 primary antibody and Alexa Fluor 488 secondary antibody to visualize the endothelial cells. Mounted samples were examined using laser scanning confocal fluorescence microscopy with a 20x/0.8 NA objective. To examine the entire ∼20 mm^2^ tissue section, automated tiling was used together with serial sectioning to acquire ∼30 fields of view and 16 z-slices representing 15 μm/field of view. In parallel, mice not receiving AF568-LDL were processed and imaged using the same microscope settings, including scan speed and laser intensity, to determine the amount of background (autofluorescence). The images represent collapsed z-stacks with the fluorescence pseudocolored using the inverted Bordeaux LUT (https://github.com/cleterrier/ChrisLUTs/blob/master/I%20Bordeaux.lut) using ImageJ (**Figure 5B**). Although fewer fields of view were acquired for these samples, 16 z-slices were captured, and integrated pixel intensity from the samples not receiving AF568-LDL served as the threshold for processing the AF568-LDL-containing samples. As illustrated in **Figure 5C**, following 30 mins of incubation, AF568-LDL is apparent in active areas of LDL transcytosis. Insets are included to demonstrate the signal above the background and the arrangement of the CD31-positive endothelial cells pseudocolored with the inverted Forest LUT. Of note, co-administration of the HDL decreased the deposition of the AF568-LDL by ∼40% (p-value = 0.02) (**Figure 5D, E**), consistent with the notion that HDL and LDL compete for binding of the transcytosis cargo receptor, SR-B1.

**Figure 5.**
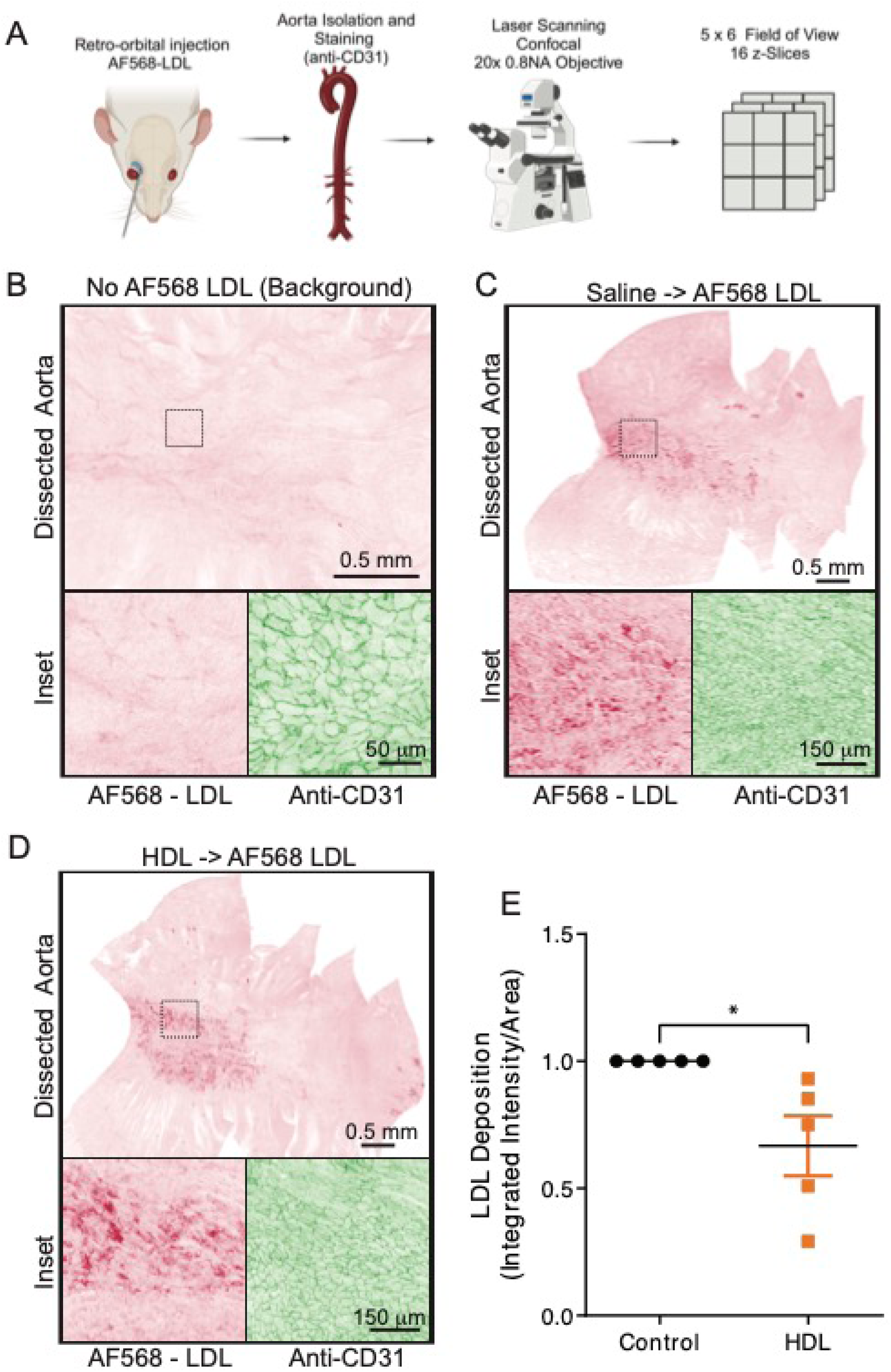
Increasing HDL levels attenuates AF568-LDL deposition in mice aorta. **(A)** Schematic outline of the *in vivo* procedure to measure LDL deposition in mice aorta from treatment to image acquisition of the isolated aorta. **(B-D)** Representative images and magnified insets of dissected and processed mice aorta, Panel B contains are from a mouse not injected with AF568-LDL, Panel C is from a mouse injected with saline and AF568-LD, and Panel D is from a mouse with human HDL and AF568-LDL co-administered. The red fluorescence of the AF568 is presented as an inverted image with the background subtracted. The black dotted box marks insets in the tiled image, and the corresponding red and green (anti-CD31) are depicted. **(E)** Quantification of the background subtracted AF568-LDL fluorescence from isolated aortas of mice receiving both HDL and AF568-LDL normalized to basal AF568-LDL deposition in PBS-perfused mice. n=5, * p<0.05.

Our pulldown experiments and MST data (**Figure 2C-E**) demonstrate that APOA1-Milano is a better ligand for SR-B1 than wild-type APOA1. However, the cell-based assays were less conclusive in distinguishing a difference between the two isoforms. Given the results from **Figure 5**, we examined whether APOA1-WT could also limit the acute deposition of AF568-LDL in the mice aorta and if APOA1-Milano had a more significant impact. To this end, the APOA1 variants were injected into mice to determine their ability to limit the deposition of injected AF568-LDL in the aorta in C57BL/6J male mice (**Figure 6**). LDL deposition was reduced by ∼30% (p-value <0.05) when a 2-fold excess (by protein mass) of wild-type APOA1 protein was perfused 30 min before the injection of the AF568-LDL LDL (Figure 6A, B, **and D**). Remarkably, perfusion of APOA1-Milano before AF568-LDL was able to block LDL deposition by about 60% (p-value < 0.0001) (**Figure 6 C, D**), indicating that APOA1-Milano is a more potent inhibitor of acute deposition of LDL in this animal model.

**Figure 6.**
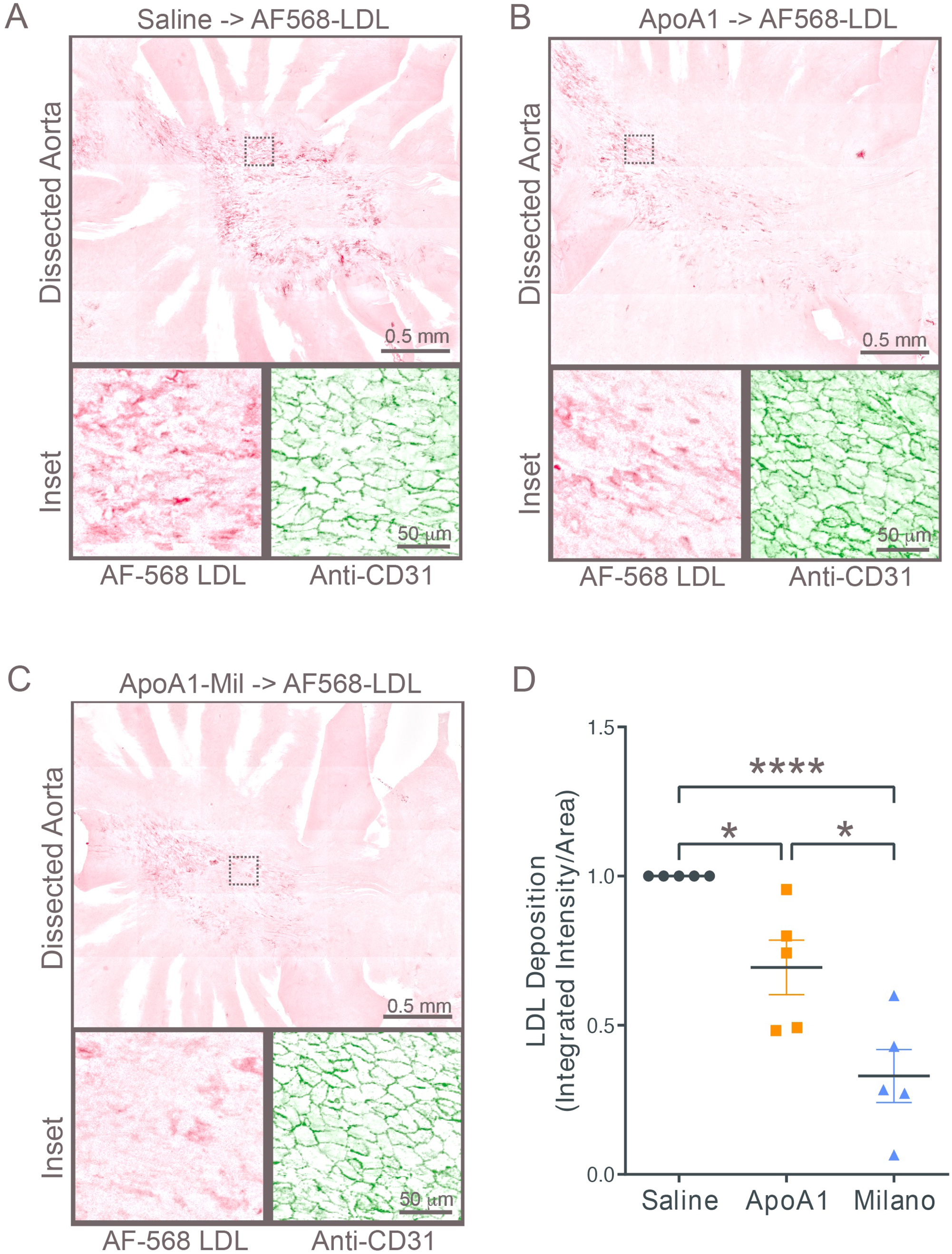
APOA1 and APOA1-Milano inhibit deposition of AF568-LDL in mice aorta. **(A-C)** Representative images of *en face* staining of the isolated aortas from mice perfused with PBS or 0.4 mg of APOA1-WT or APOA1-Milano prior to perfusion with AF568-LDL through retro-orbital injection. **(D)** Quantification of background corrected AF568-LDL fluorescence from *en face* staining of the isolated aortas; values are normalized to basal AF568-LDL deposition in PBS-perfused mice. n=5, * p<0.05, **** p<0.0001.

## Discussion

Using high-resolution microscopy and fluorescently labeled lipoproteins, the results from this study suggest that circulating HDL can slow the development of cardiovascular disease by limiting LDL transcytosis and deposition in the aorta via competitive associations with SR-B1. Data from previous studies using non-physiological concentrations of HDL support this general notion.^4,65^ Our cell-based assays used physiological concentrations of rHDL2, described as the preferred ligand for SR-B1,^60^ demonstrating that LDL uptake and transcytosis can be partly inhibited.

HDL is cardioprotective through several potential mechanisms, and this study reveals yet another contributing mechanism. HDL is well known to accept cholesterol for reverse cholesterol transport. Additionally, HDL is also an anti-inflammatory particle^9–11^ and known to carry proteins such as paraoxonase 1 (Pon1) and platelet-activating factor acetyl hydrolase (PAF-AH) that are anti-oxidative proteins.^12–15^ HDL has been shown to induce the production of the vasodilator prostacyclin,^21^ promotes endothelial integrity by inhibiting oxidized LDL-induced apoptosis^18,19^ and through SR-B1, stimulates endothelial cell migration for wound healing^20^ and nitric oxide production.^16^ Furthermore, HDL also carries sphingosine-1-phosphate (S1P),^16^ a potent anti-apoptotic sphingolipid, which may contribute to the anti-apoptotic capabilities of HDL.^17^ Finally, HDL also carries miRNAs, such as miR-126 and miR92a^22^ which are involved in vascular biology and inflammation, and miR-223, which regulates SR-B1 expression.^23^ The importance of the individual processes and their relative contribution to the prevention of cardiovascular disease remains unknown.

Carriers of the Milano mutation have low levels of plasma HDL and elevated triglycerides but do not present any features of HDL deficiency disorders and show no evidence of cholesterol deposition in tissues or atherosclerotic vascular diseases.^49,52^ Understanding the mechanistic changes that explain why Milano carriers have low APOA1 levels yet are resistant to cardiovascular disease has been elusive. The APOA1-Milano mutation, Arg163 to Cys163, provides a potential intermolecular disulfide bond formation and dimerization of two APOA1-Milano proteins. One possibility is that the dimers have improved functionality over the monomers. From a structural perspective, the arginine in APOA1-WT forms an intrahelical salt bridge with glutamate at position 169; this would be lost in the Milano mutation.^66^ It is conceivable that loss of the salt bridge would result in APOA1-Milano being less stable and having more exposed hydrophobic regions than APOA1-WT.^66^ However, direct examination of the secondary structure of APOA1-Milano has led to conflicting conclusions, with studies observing an increase in alpha-helical content,^67,68^ while other studies report decreases^66,69^ or even no change compared to APOA1-WT.^70^ Reconstituted HDL^Mil^ had a superior ability to inhibit LDL uptake compared to the free APOA1-Milano (**Figure 3B, 4B**). This suggests that the interaction of the APOA1 isoforms with SR-B1 remains intact and that the arrangement of APOA1-Milano and APOA1-WT in the HDL particle is not grossly impacted. Indeed, the structure of wild-type and Milano APOA1 within HDL is thought to be similar based on crosslinking and mass-spectrometry studies.^71,72^ Further structural studies of APOA1-Milano (full length or truncations) through X-ray crystallography or NMR would provide more insight into potential structural differences between wild-type and Milano APOA1.

Cardiovascular disease remains a source of morbidity and mortality without a cure. Considering the observation that ALK1- and SR-B1-mediated LDL transcytosis are necessary for the formation of atherosclerosis, we propose that an additional mechanism by which APOA1 and HDL are beneficial is by the continuous competitive binding to SR-B1. By antagonizing LDL uptake and, ultimately transcytosis to the intimal space, HDL will help slow the development of atherosclerosis. Contextually, our results also suggest that small molecules or inhibitors that block LDL transcytosis may be beneficial in limiting cardiovascular disease. However, given the role of SR-B1 in reverse cholesterol transport to hepatocytes for bile acid production and steroidogenic cells, we suspect that targeting Alk1 may be a more prudent course of action.

## Acknowledgments

The authors would like to thank the Kennan Research Center for Biomedical Science Core Facilities at St. Michael’s Hospital, especially Dr. Caterina Di Ciano-Oliveira and Dr. Monika Lodyga, for their continued support, technical advice, expertise, and training.

## Data and materials availability

All data is available in the main text or supplementary materials.

## Supplemental Figure Legends

**Supplemental Figure 1. SDS PAGE analysis of the purification of the APOA1 variants. (A)** SDS PAGE analysis of the purification process of the three APOA1 variants where the arrow indicates the APOA1 protein. The loading order is of the following: for APOA1-WT and APOA1-Monomer, 1) Not Induced, soluble 2) 1 mM IPTG induced, soluble, 3) Flow thru from purification, 4) Wash 4 (10 mM Imidazole), 5) Wash 5 (10 mM Imidazole), 6) Elution with 0.1 M EDTA, 7) Dialyzed Elution (in PBS). For APOA1-Mil: 1) Not Induced, soluble 2) 1 mM IPTG induced, soluble, 3) Wash 4 (10 mM Imidazole), 4) Wash 5 (10 mM Imidazole), 5) Elution with 0.1 M EDTA, 6) Dialyzed Elution (in PBS). **(B)** The resulting SDS PAGE gel of samples treated with or without DTT to assess the disulfide bond state of APOA1-Mil. The loading order is of the following: 1) APOA1-WT 2) APOA1-Mil 3) APOA1-Monomer.

**Supplemental Figure 2. Heterogeneity in the amount of DiI-LDL internalized in human coronary endothelial cells as measured through DiI fluorescence per cell. (A)** Representative image of DiI-LDL internalization in HCAECs accompanied with **(B)** quantification of the integrated density of DiI-LDL fluorescence per cell of a typical replicate experiment.

**Supplemental Figure 3. Assessing the generated rHDL^APOA1^ particles. (A)** Native PAGE (no SDS and non-reducing) analysis to gauge the size of the generated DiI-rHDL^APOA1^ (lane 4) made from recombinant APOA1 (lane 3) in comparison to isolated DiI-LDL (lane 1) and commercially purchased HDL (lane 2). The gel was generated with made with polyacrylamide and the system ran at pH 8.75. **(B)** Plot of the density of the generated DiI-DMPC-WT particles measured manually along with absorbances at 280 nm and 555 nm to detect APOA1-WT and DiI respectively).

**Supplemental Figure 4. Verifying bi-cistronic SR-B1 + GFP plasmid through IF and western blot and association specific association with APOA1 containing particles. (A)** Representative images of HeLa cells transfected with GFP alone or the bi-cistronic SR-B1 + GFP plasmid stained with a Cy3-conjugated SR-B1 nanobody. **(B)** Lysate collected from HeLa cells transfected with GFP alone or the bi-cistronic SR-B1 + GFP plasmid ran on a western blot and was probed with an SR-B1 antibody to verify that full-length SR-B1 was overexpressed compared to the GFP-alone transfected HeLa cells. **(C)** Quantification of uptake of different densities (all of which span the density of natural HDL) of DMPC and DiI particles that lack APOA1 by SR-B1 + GFP and GFP overexpressing HeLa cells. n=4 for <1.08 g/mL, n=3 for 1.08-1.09 g/mL and for >1.1 g/mL, where 15 cells were imaged and analyzed per condition per n.

