## Supplementary figures and images for "Apolipoprotein A1 and high-density lipoprotein limit low-density lipoprotein transcytosis by binding scavenger receptor B1"

### supplemental figures

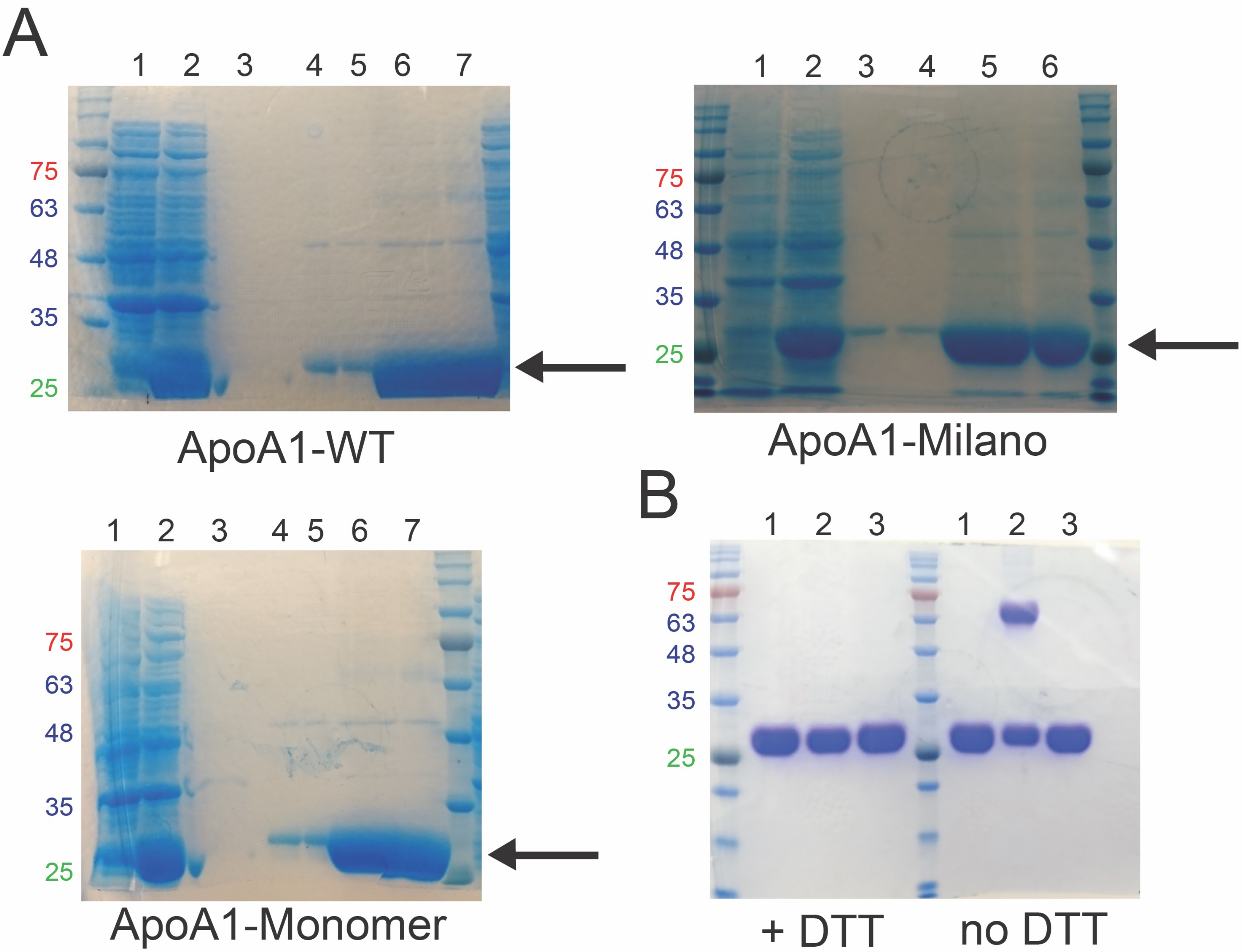

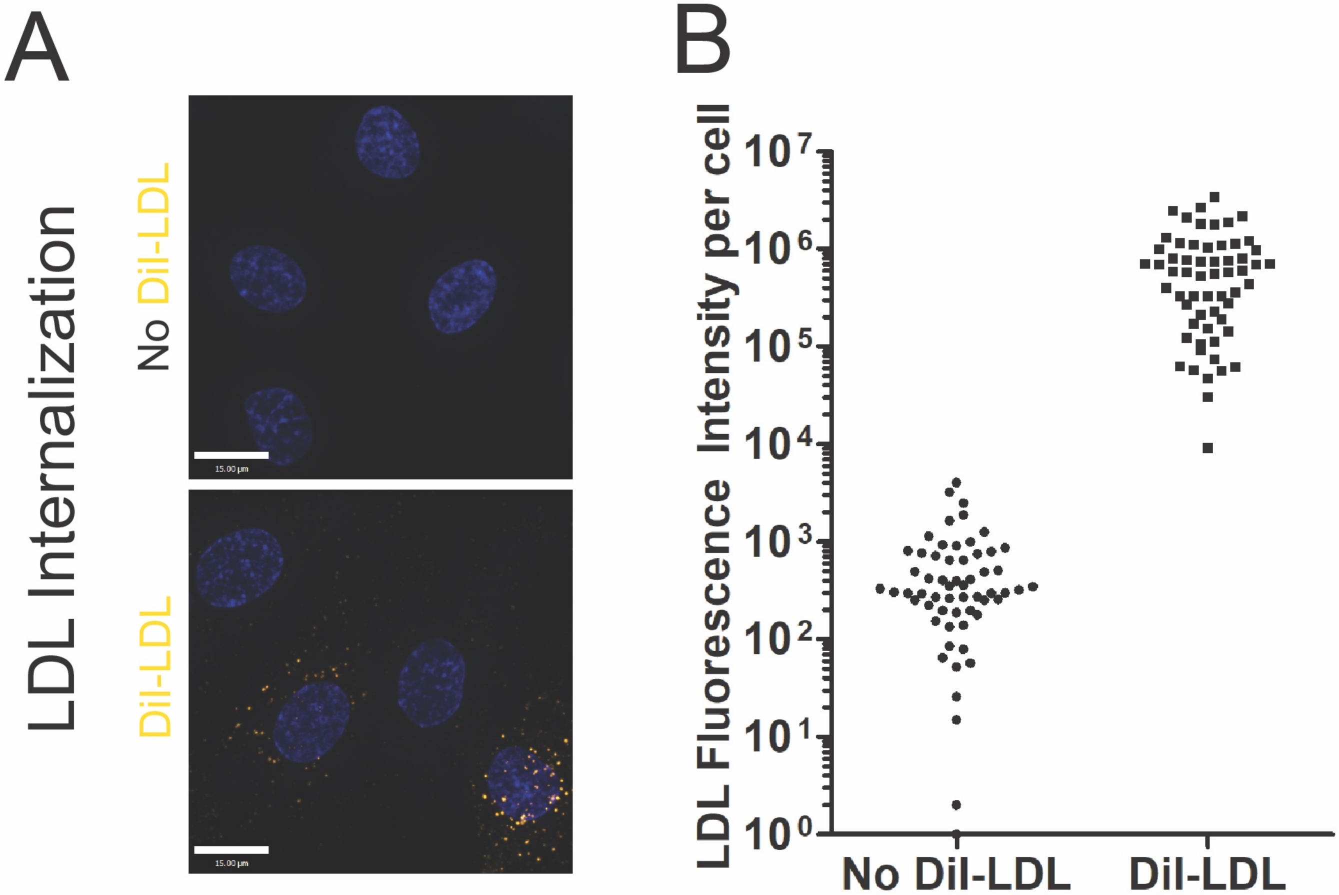

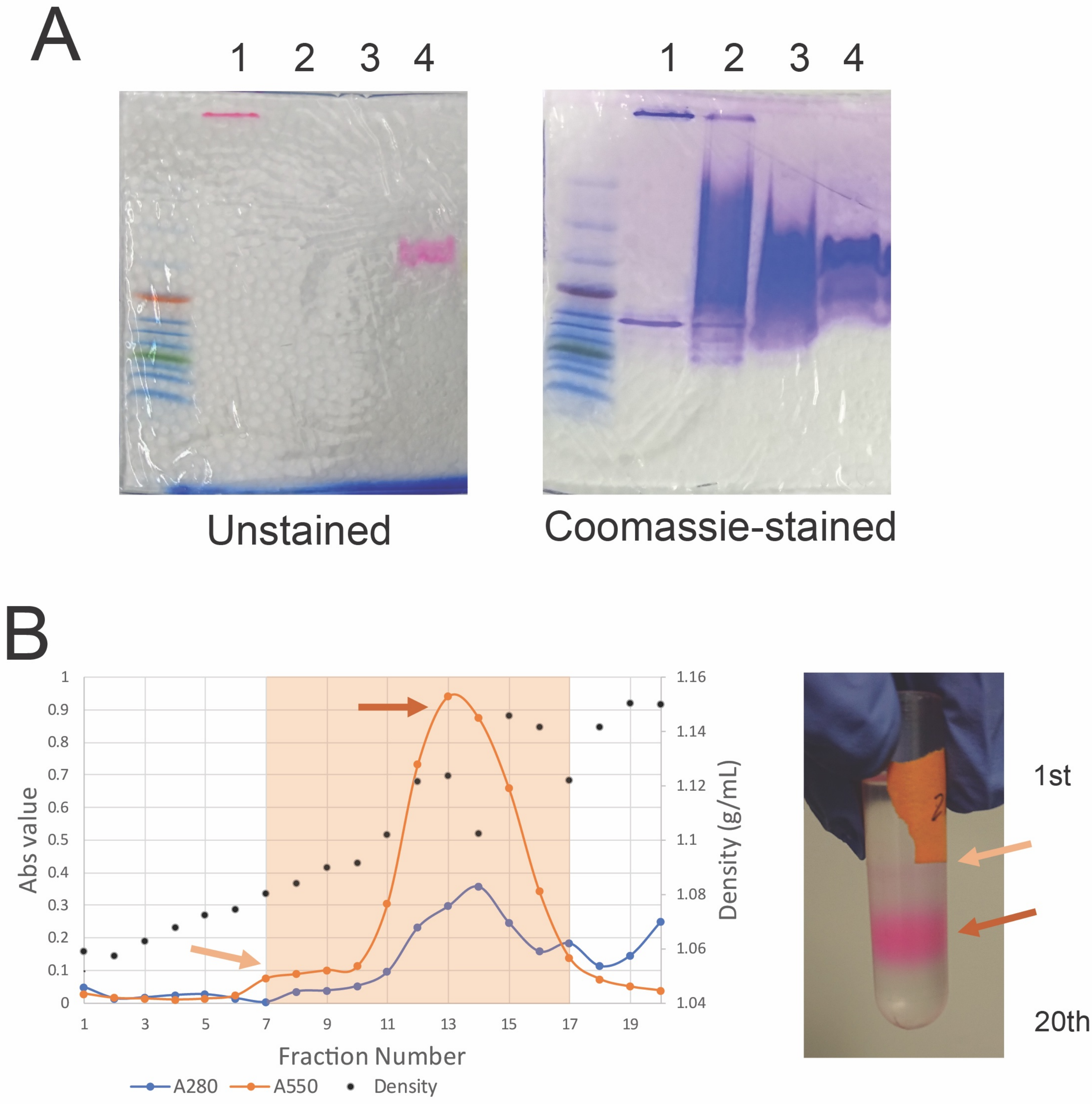

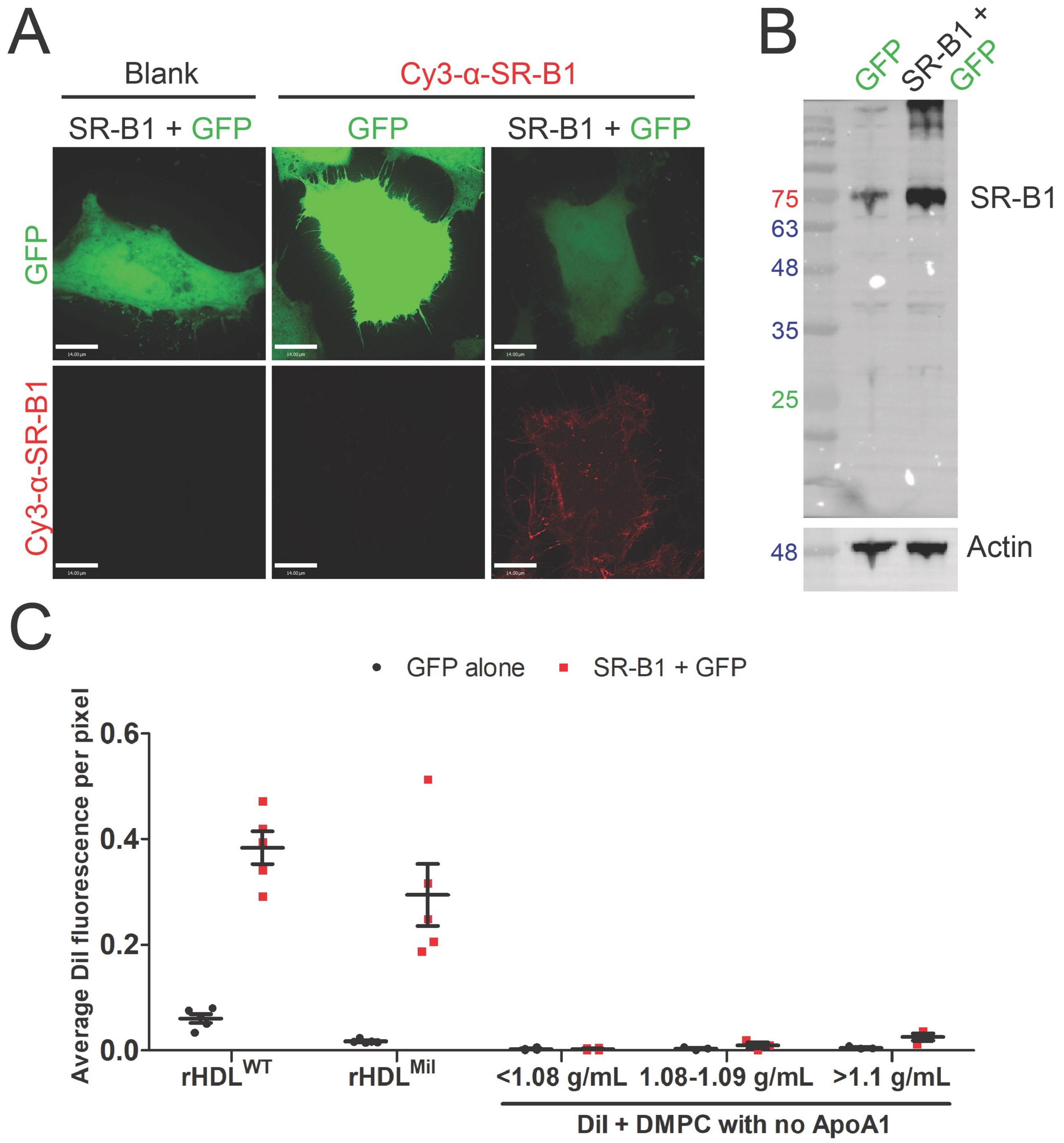
